# Admixture mapping among three hybridizing sapsucker (*Sphyrapicus*) species shows importance of the Z chromosome in phenotypic differentiation and speciation

**DOI:** 10.64898/2026.08.15.745054

**Authors:** Libby Natola, Jocelyn Hudon, Darren Irwin

## Abstract

Plumage pigmentation is under intense sexual and natural selection and plays an important role in the speciation process in birds, so there is much interest in uncovering the genomic basis of plumage colour differences between populations and species. Three species of North American woodpeckers, the red-breasted (*Sphyrapicus ruber*), red-naped (*S. nuchalis*), and yellow-bellied sapsuckers (*S. varius*), provide a particularly promising opportunity to unravel the genomic mechanisms of plumage colour differentiation. The breeding ranges of the three species are mostly non-overlapping but adjacent, with hybrid zones occurring where the ranges meet. The species pair with the most similar plumage colouration (*S. varius* and *S. nuchalis*) is not the most closely related pair genomically (*S. nuchalis* and *S. ruber* is), providing an opportunity to determine the subset of the genome that underlies the plumage colour variation. Using admixture mapping of whole genome sequences from each species and hybrids from each species pair, we show close associations between the colour of multiple plumage patches and wide swathes of the Z-chromosome. These results highlight how comparable changes in one sex chromosome can cause either slight plumage pigmentation changes (*S. varius* vs. *S. nuchalis*) or large-scale shifts from dimorphism to monomorphism and from primarily black and white to primarily red plumage colouration (*S. varius* and *S. nuchalis* vs. *S. ruber*).

## Introduction

For centuries evolutionary biologists have attempted to explain the diversity of integumentary colouration in birds and its relationship to speciation and taxonomy (Mayr, 1942; Wallace, 1877). Plumage colouration plays an important role in mate attraction and choice (Baker & Boylan, 1999), territorial displays (Pryke et al., 2001), quality signaling (Weaver et al., 2018), and predator evasion (Gluckman & Cardoso, 2010), and is therefore under intense sexual and natural selection pressures. However, plumage colouration is a complex trait that involves developmental pathways controlling melanin production, carotenoid deposition, and feather structure, complicating attempts to infer the physiological and molecular bases of plumage colour variation (Cuthill et al., 2017; Funk & Taylor, 2019).

More recently, ornithologists have applied advanced genomic methods (Baiz et al., 2020; Funk & Taylor, 2019) to better understand how and why so much plumage colour diversity exists. The advent of next generation sequencing methods has yielded genetic associations with a variety of plumage colours and patterns. While in some cases, evidence points to plumage colour modifications being caused by small changes in a single protein-coding gene (Theron et al., 2001) or in a regulatory region (Gazda et al., 2020; Lopes et al., 2016; Toews et al., 2016), colour transitions have also been linked to large chromosomal inversions (Huynh et al., 2011). Variation in plumage colour traits may be caused by a few genes with large effect (Toews et al., 2016) or many genes with small effect (Aguillon et al., 2021). Sometimes the same genes are involved across taxa, as seen with ASIP (Stryjewski & Sorenson, 2017; Toews et al., 2016; Wang et al., 2020; Whitlock, 2015) and BCO2 (Gazda et al., 2020; Toews et al., 2016), while different pathways achieve similar changes in other species (e.g., RALY, Wang et al., 2020). The genomic changes affecting these phenotypes can arise through apparently novel mutations (Huynh et al., 2011; Theron et al., 2001) or via introgression from related species (Baiz et al., 2020; Gazda et al., 2020).

Three North American woodpecker species, the red-breasted, red-naped, and yellow-bellied sapsuckers, began to differentiate approximately one million years ago (Weir & Schluter, 2004) but these three forms hybridise in pairs or in a triad in regions where their breeding ranges overlap (Billerman et al., 2016, 2019; Grossen et al., 2016; Howell, 1952; Johnson & Johnson, 1985; Natola et al., 2022; Scott et al., 1976; Seneviratne et al., 2012, 2016). All have unique plumage colour patterns (Figure 1), the genetic basis of which has been theorised since Johnson and Johnson (1985) speculated the existence of six gene pairs affecting plumage colouration in red-breasted and red-naped sapsuckers. However, the technology to identify these loci did not exist until recently. Using Genotyping-by-Sequencing and association mapping, Grossen et al. (2016) identified a single SNP locus on the COG4 gene that is associated with plumage colour variation among the three species of sapsuckers and their hybrids, which they suggested may be a duplicated gene. In sapsuckers, the plumage colour patterns belie their phylogenetic relationships, as the most closely related species are not most similar in plumage colouration (Figure 1). The red-breasted sapsucker has an exceptionally divergent plumage from the other two sapsuckers and is also the only sexually monochromatic form. Yet the red-breasted sapsucker and the red-naped sapsucker are much more closely related genetically to each other than either is to the yellow-bellied sapsucker, which looks similar to the red-naped sapsucker. This difference in relationships between the molecular genetics and plumage colouration makes these species excellent candidates to study the genetic determination of plumage traits by allowing the decoupling of trait similarity from shared ancestry.

**Figure 1.**
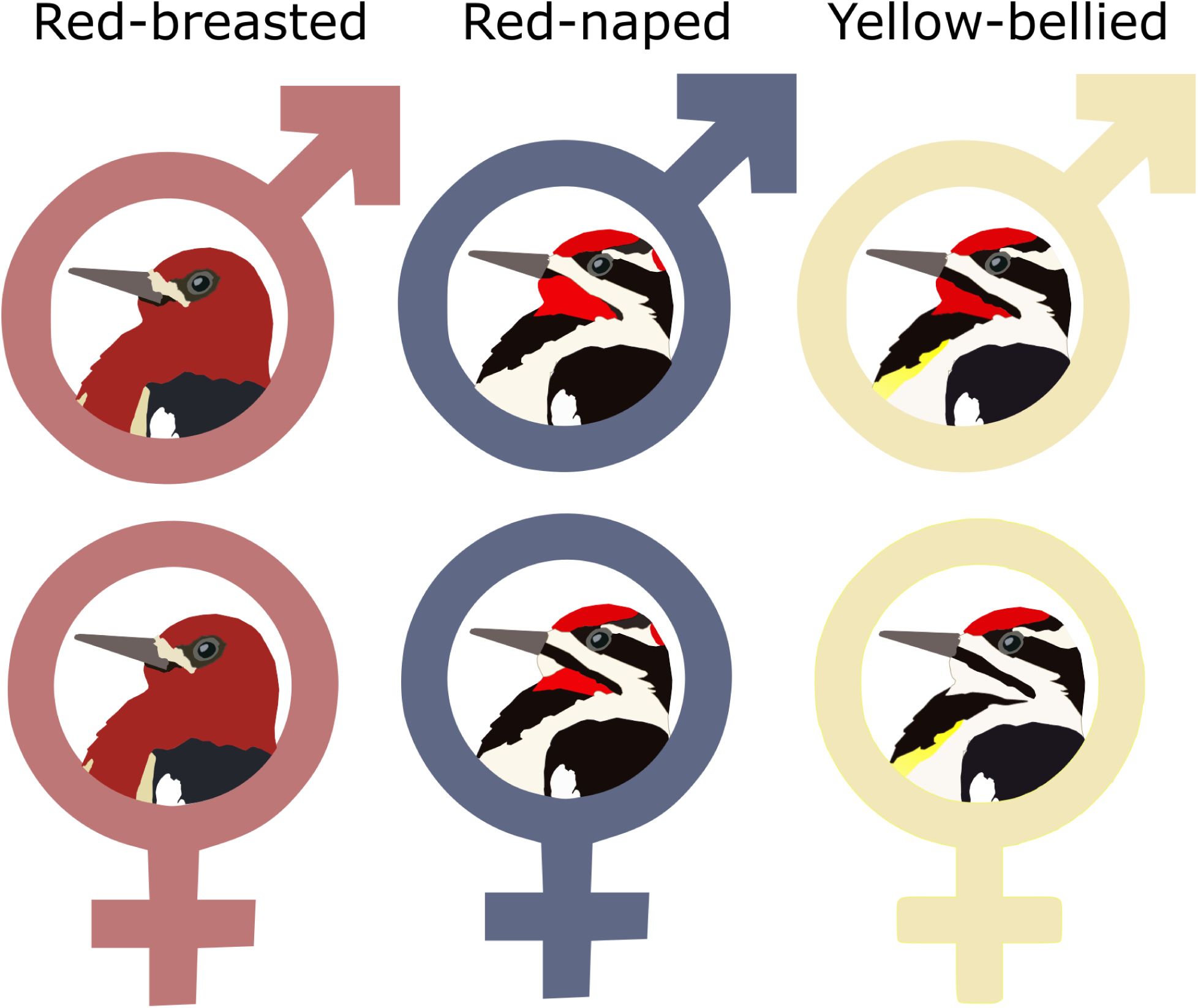
Depictions of typical parental plumage patterns for each sex of each species. Red-breasted sapsuckers are sexually monochromatic.

Identifying the mechanisms of sapsucker plumage colour differentiation is a step towards better understanding the role of plumage colour in evolution. The clear differences in plumage colouration may promote assortative mating, a process which would limit hybridization between the different species. However, some differences could lead to asymmetrical hybridization as well. In a study of breeding pairs of sapsuckers in a hybrid zone, assortative mating did predominate, with only 6.2% of mated pairs being interspecific (Johnson & Johnson, 1985). Yet, eight of the nine interspecific nests consisted of a red-breasted male mated to a red-naped female, suggesting that female sapsuckers may be more likely to mate with another species if it is redder (Johnson & Johnson, 1985). Preference for red colouration has been shown to be common in avian taxa (Plenge et al., 2000; Qvarnström et al., 2004; Ranasinghe et al., 2024), and it is reasonable to expect sapsuckers to behave similarly. It is unclear how plumage colouration operates in sapsucker mate choice, but it is likely important.

Here, we use admixture mapping analysis of whole genome shotgun resequencing (WGS) and plumage colour data for all three sapsucker species and their hybrids, to identify the genomic loci and regions underlying plumage colour differentiation in this superspecies. We are particularly interested in whether loci associated with plumage colour differences in other systems will also be implicated in this system, and whether plumage transitions are due to many genes of small effect or a few genes of large effect. Based on preliminary genotyping-by-sequencing data (GBS) (Grossen et al., 2016), we expect to see a duplication event of the COG4 gene associated with red-breasted sapsucker plumage. However, this research goes beyond the work of Grossen et al. (2016) in that we sampled from all three hybrid zones (Grossen et al. sampled only red-breasted × red-naped and red-breasted × yellow-bellied zones) using Whole Genome Shotgun sequencing data, which achieves a much denser sampling of the genome than GBS. In addition, we expect to find that plumage colouration in these species is dictated by few genes of large effect based on mis-matches between phenotypic and genotypic species assignments using GBS data, particularly in yellow-bellied sapsuckers, documented in previous studies (Natola et al., 2022; Natola & Irwin, 2025).

## Materials and Methods

### Plumage data collection

We measured associations between plumage phenotypes and genotypes by collecting plumage and genotypic data for 76 sapsuckers: 30 vouchered museum specimens from allopatric populations of each species (10 per species); 45 from field collected data within the central BC red-breasted × red-naped hybrid zone, the northern BC red-breasted × yellow-bellied zone, and the west-central AB yellow-bellied × red-naped zone (15 from each hybrid zone); and one wild-caught tri-species hybrid (Figure 2). All samples included here were used in Natola and Irwin (2025), and some were also used in Natola et al. (2023), Natola et al. (2022), Seneviratne et al. (2016), and Grossen et al. (2016). To maximise the statistical power to detect differences on the W chromosome (found only in females), we chose to weight our sampling towards females from allopatric populations (∼60% female by field identification). In order to maximise detection of genes underlying variation in plumage colouration (which varied most in males), we weighted our sampling towards males from sympatric populations (∼66% male by field identification). This strategy also enabled us to investigate patterns of sexual dichromatism versus monochromatism. We chose 12 variable plumage patch locations to measure among all species including the back, black nape, white nape, crown, above superciliary, superciliary, below superciliary, auricular, below auricular (sometimes referred as the malar stripe), chin, throat, and breast band (Figure 3).

**Figure 2.**
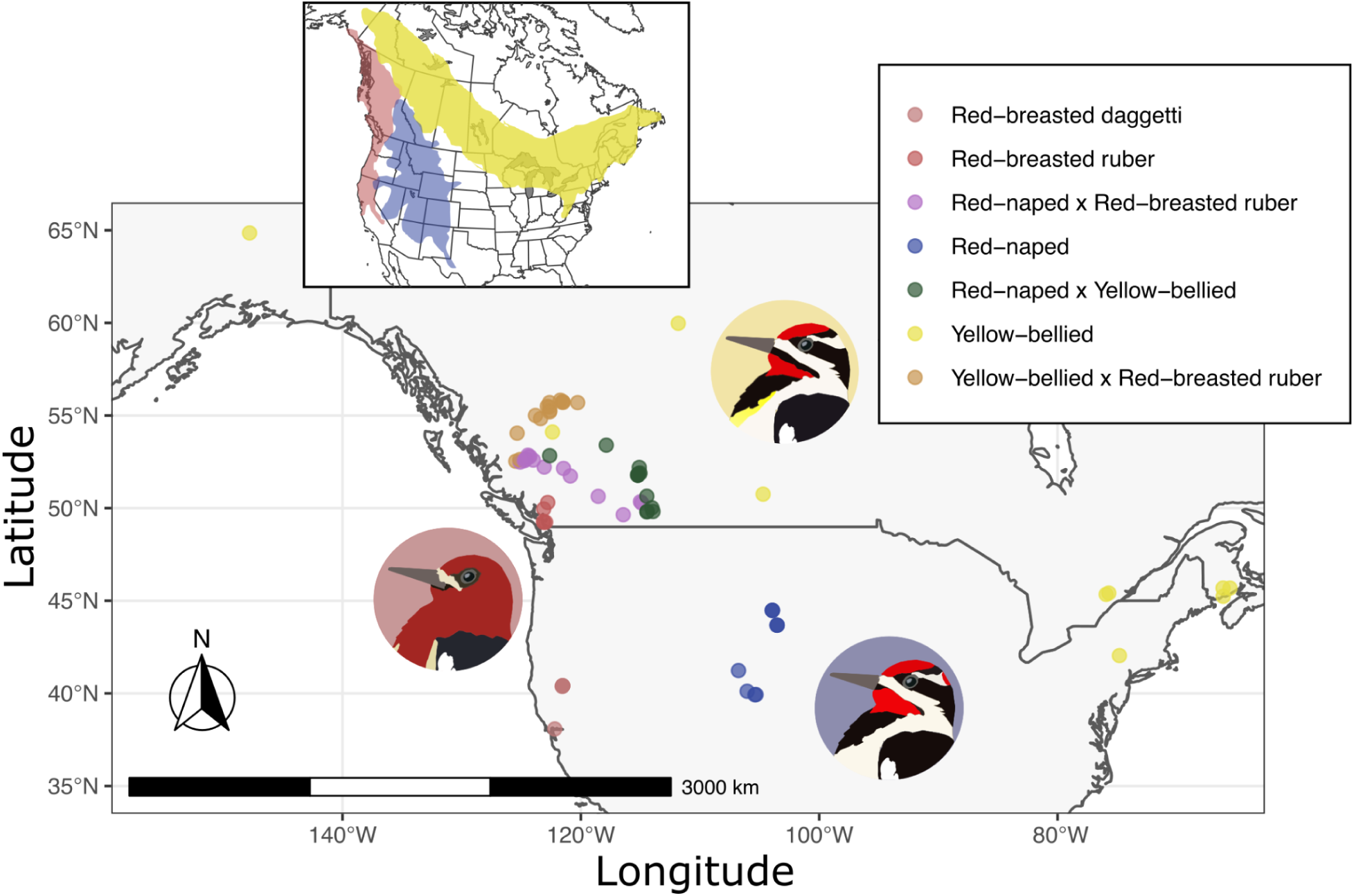
Map of sampling locations for this study, adapted from Natola and Irwin 2025.

**Figure 3.**
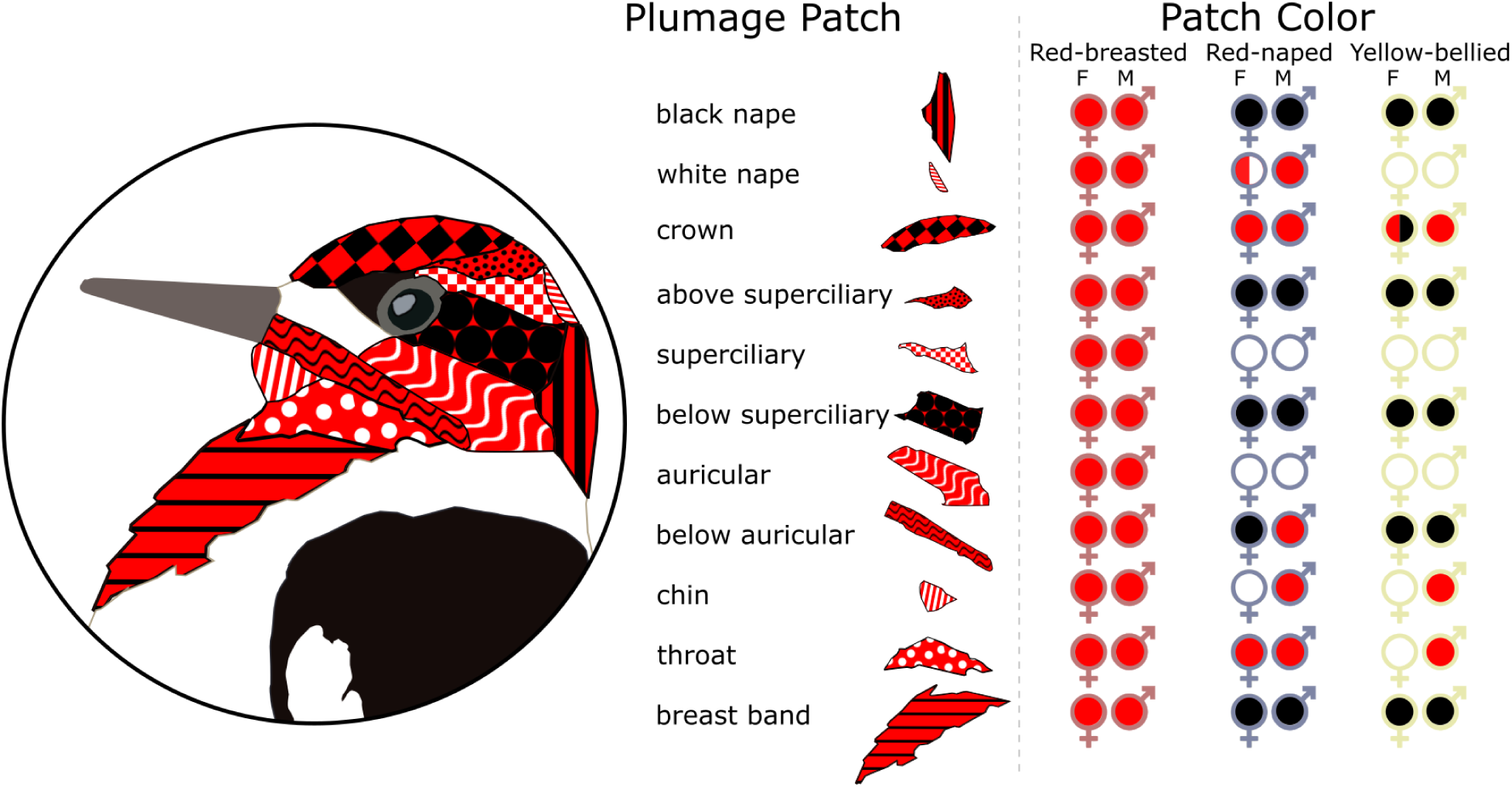
Plumage patches scored in this study. Pattern colours represent colours seen in any plumage patch (red, black, and/or white). Patch colour section indicates sex and species difference for each patch with the symbol outline colour indicating species, and the fill indicating the colour of each patch in the circle for each sex and species.

### Genotypic data collection

See Natola and Irwin (2025) for field and museum tissue collection, Whole Genome Shotgun (WGS) Sequencing, and genotypic data processing methods. In brief, we extracted DNA from 76 samples (10 allopatric birds from each species, 15 from each hybrid zone representing a range of admixture proportions, and one putative three-way hybrid), with a standard phenol-chloroform protocol. We submitted 100 ng DNA to Genome Quebec for library preparation using the NEB Ultra II (New England Biolabs) library preparation kit and sequencing on four Illumina NovaSeq6000 S4 PE150 lanes. WGS reads were filtered and aligned to a red-breasted sapsucker reference genome (Natola & Irwin, 2025) assembled *de novo* from Pacific Biosciences SMRT long reads, scaffolded to a golden-fronted woodpecker genome (Wiley & Miller, 2020), and annotated by mapping sequences to the zebra finch genome (Rhie et al., 2021) with the program liftoff. Using bamqc outputs from Qualimap v 2.2.2a (Okonechnikov et al., 2016), we plotted WGS coverage of each individual bird’s mapped read coverage for each scaffold divided by the bird’s genome-wide mean coverage for a corrected coverage measure. We then manually checked histograms of corrected coverage. Any scaffolds with corrected coverage of approximately one in known males and 0.5 in known females were designated likely Z chromosome scaffolds and scaffolds with corrected coverage of approximately zero in males and 0.5 in females were presumed W chromosome scaffolds (See Natola and Irwin, 2025).

## Analysis

### Linear Mixed Models

To prepare reads for analysis in GEMMA v. 0.98.3 (Zhou & Stephens, 2012), we removed two red-naped samples with insufficient phenotypic data, phased and imputed genotypes using Beagle v. 5.2 (Browning & Browning, 2007), then converted to .ped format using PLINK v. 1.9 (Purcell et al., 2007). We used GEMMA to generate a relatedness matrix for all samples. We ran separate Univariate Linear Mixed Models (LMM) for each trait in GEMMA, with sex as a covariate. To determine where to draw our significance thresholds for each trait, we repeatedly randomised the phenotypes for hybrid-zone birds separately for males and females and ran this randomised GEMMA for each trait. We did this 100 times per trait, then retained both the highest and the 100^th^ highest -log(*p*) values (i.e., the lowest and 100^th^ ranked lowest *p* values) from all SNP associations across all the randomizations for each trait. With ∼ 10 million SNPs run in 100 randomizations, under the null hypothesis of no real association the probability of surpassing the high threshold for any SNP is approximately 1 in one billion, and surpassing the lower threshold is 1 in 10 million. Any SNP falling above the highest SNP threshold is likely truly associated with the trait, but we might expect 1 unrelated SNP to occur above each trait’s lower threshold per LMM. We therefore have high confidence in any associations with a -log(*p*) above this threshold, and SNPs for each trait were considered outliers if their likelihood surpassed the trait’s highest random association threshold. For all traits that varied within or between sexes, we ran LMMs with sexes separated. We ran males only for traits which were variable among males of the different species (white nape, below auricular) and females only for traits that varied among females of each species (white nape, below auricular, chin, throat). Though crown colour varies among yellow-bellied sapsucker females (from almost completely black, to having white spots, or being completely red), there was not enough variation in phenotypes of our study individuals to run the LMM for the crown of females alone. We also ran Bayesian Sparse Linear Mixed Models to identify those SNPs with the highest likelihood of involvement in the trait; see Supplemental Materials for details and results (Figure S1).

### Locus selection, gene ontology analysis

For each SNP associated with a phenotype via LMM (-log(*p*) > the trait randomization cutoff) that was not on a Z-chromosome scaffold, we used BEDtools v 2.30 (Quinlan & Hall, 2010) to compile a list of the closest genes both up- and downstream from the lifted-over gene annotation (based on a zebra finch annotation) of the red-breasted sapsucker reference genome (Natola et al., 2022). We then searched for information regarding these genes in the UniProt database.

Because several thousand SNPs on the Z chromosome scaffolds had high associations with plumage traits, we were unable to identify specific putative causal genes for these traits. Instead, we compiled a list of all characterised genes mapping to the Z-chromosome scaffolds. To see if any matched known pigmentation or colouration genes, we cross-referenced the list of Z-chromosome mapped genes to the curated lists on IFPCS (International Federation of Pigment Cell Societies, ifpcs.org/colourgenes/) and GePheBase (gephebase.org). We also cross referenced genes found on the Z chromosome that have been identified as playing a role in pigmentation in other recent publications of avian colour (Aguillon et al., 2021; Baiz et al., 2020; Funk & Taylor, 2019; Lopes et al., 2016; Toews et al., 2016; Wang et al., 2020). These lists tend to be dominated by genes known to control melanin pigmentation.

## Genotype-by-individual pl2ots

To identify haploblocks associated with genotypes, we filtered loci to only those associated with a particular trait via LMM (-log(*p*) > trait randomization cutoff), then plotted genotypes for each locus for each individual using scripts adapted from Irwin et al. (2016).

### COG4 Locus

Grossen et al. (2016) found a SNP within the COG4 gene was highly associated with sapsucker plumage types. Neither the COG4 gene nor any of its aliases (COD1, CDG2J, SWILS) was included in the liftoff gene annotation. To find its location in our assembly, we selected the 80 bp surrounding the SNP position (chr11, 5,606,038 bp) in the downy woodpecker genome (*Dryobates pubescens*; Zhang et al., 2014), which Grossen et al. (2016) used to align their reads.

## Results

### Linear Mixed Models

The LMMs identify two major patterns within the sapsuckers. Six of the traits show extremely strong association with many SNPs on the Z chromosome (-log(*p*) > 25). These include the black nape, above superciliary stripe, superciliary stripe, below superciliary stripe, auricular stripe, throat, and breast band (Figure 4). Only the throat patch shows peaks of several close loci with associations surpassing our significance threshold in autosomal regions. In contrast, regions of high association with plumage patches on the sex chromosomes are so numerous that it is not possible to identify any causal SNPs as they are in tight linkage in a highly variable region. We refer to these as the “Z chromosome haploblock” traits. Notably, these Z chromosome haploblock traits are almost exclusively plumage patches that distinguish red-breasted sapsuckers from both red-naped and yellow-bellied sapsuckers (except throat).

**Figure 4.**
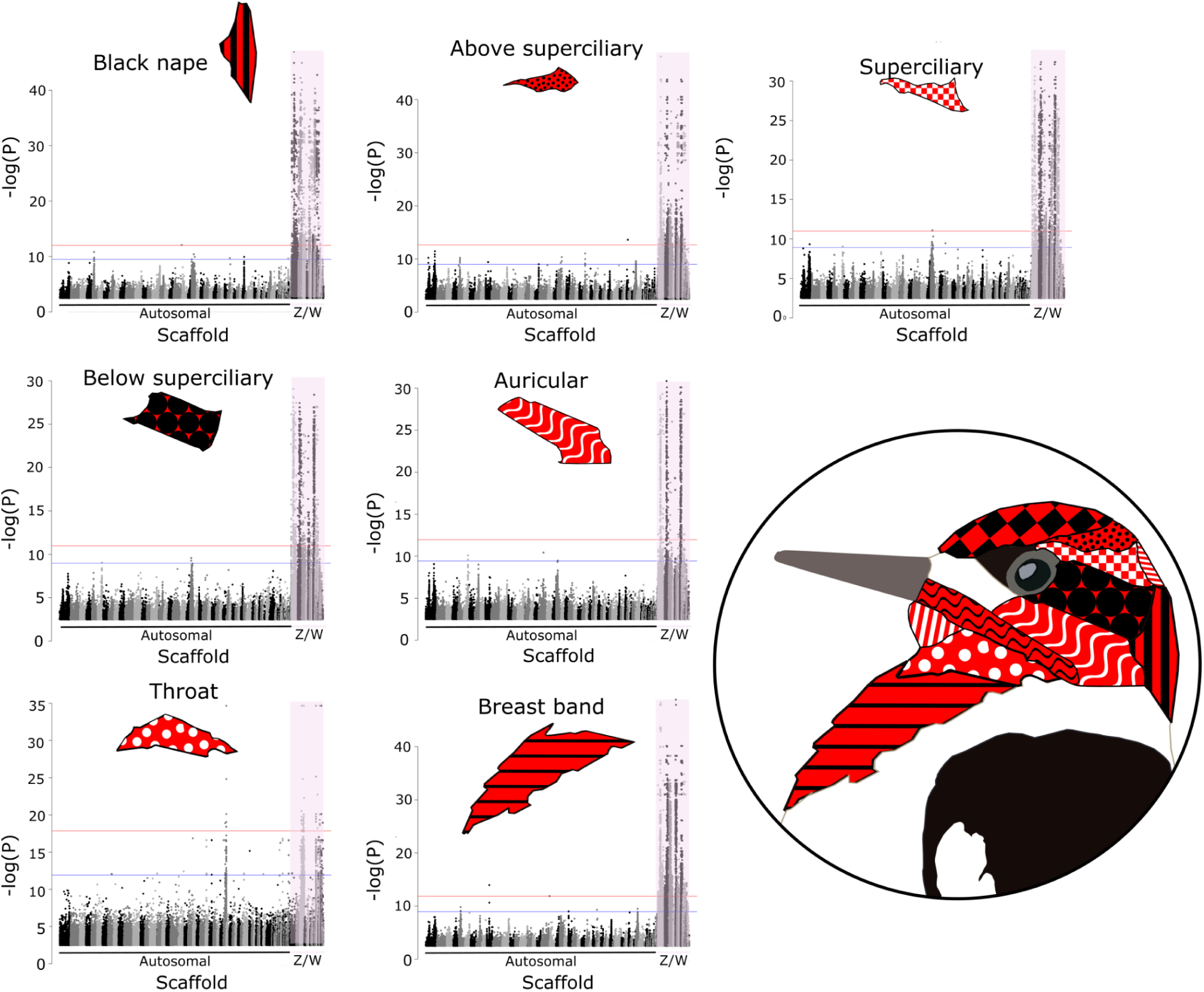
Plots showing strength of association between each SNP and a phenotypic trait, as estimated using LMM output from GEMMA for the Z chromosome haploblock traits and include all species and species crosses as well as both sexes. From left to right, black nape, above superciliary stripe, superciliary stripe, below superciliary stripe, auricular stripe, throat, breast band. The blue line indicates the 100th highest randomised -log*(p)* value threshold and the red line indicates the highest randomised -log*(p)* value threshold. Sex chromosomes (Z/W) are indicated by violet shading.

The other plumage patches do not show as clear an association with specific genomic regions, so we call these “non-haploblock” traits. While some traits in this group show some association with loci on the Z chromosome, those loci do not span across large genomic distances or show as high -log(*p*) values. Traits following this pattern include the white nape patch, crown, below auricular stripe, and chin (Figure 5). Running sexes separately for white nape, below auricular, chin, and throat, did not resolve these associations better than running them together, likely due to small sample sizes. None of these traits are diagnostic of red-breasted sapsuckers vs red-naped and yellow-bellied, rather, most differentiate red-naped and yellow-bellied species and sexes. LMMs do not identify any loci above the 100th highest randomised -log(*p*) value threshold for the plumage colouration on the back so this trait is omitted from further analyses.

**Figure 5.**
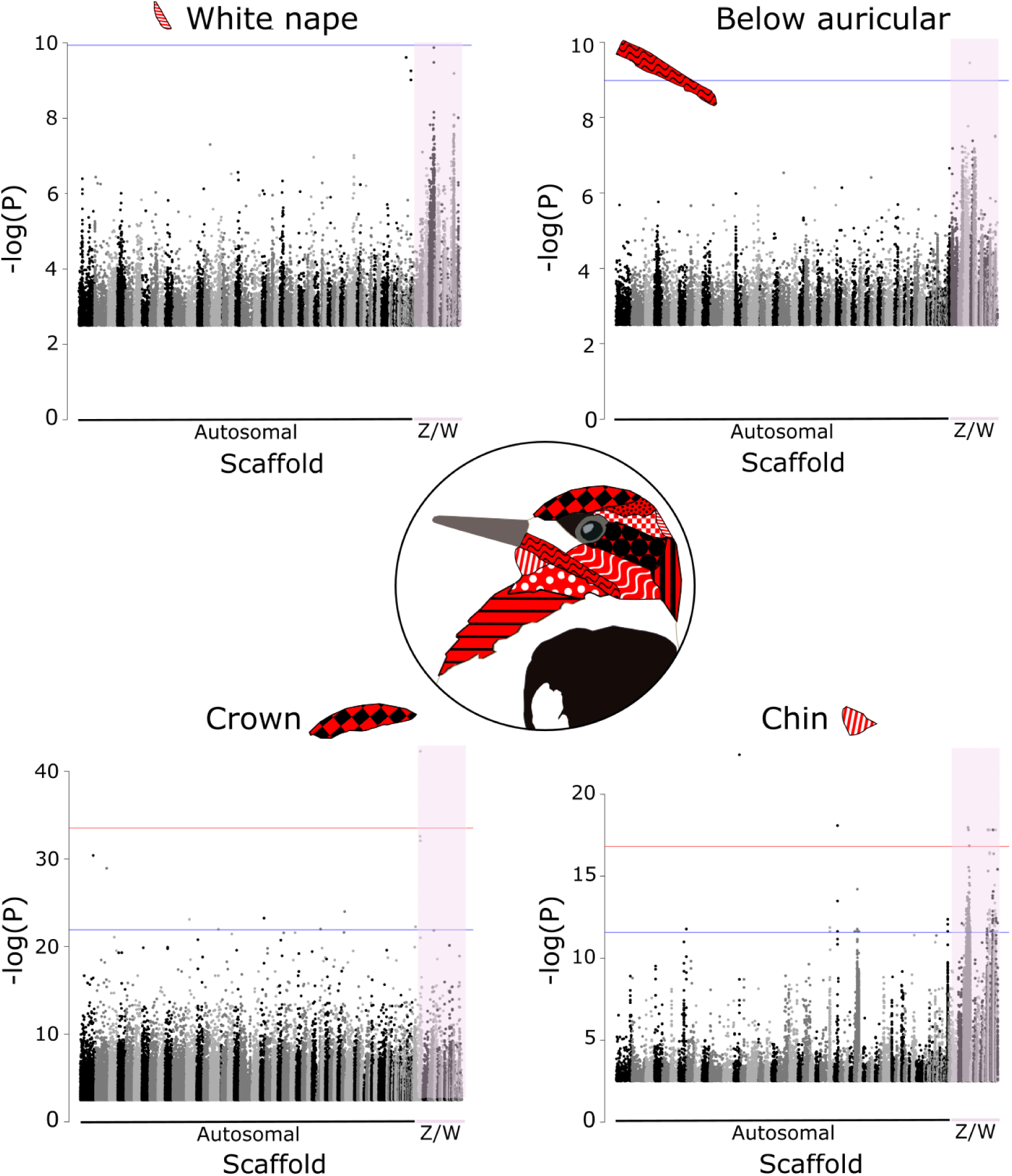
Plots showing strength of association between each SNP and a phenotypic trait, as estimated using LMM output from GEMMA in non-haploblock traits, data include all species and species crosses as well as both sexes. Plots included show output for traits in the autosome-plumage linked group. From left to right, white nape, below auricular, crown, and chin. The blue line indicates the 100th highest randomised -log(*p*) value threshold and the red line indicates the highest randomised -log(*p*) value threshold. Sex chromosomes are indicated by violet shading.

### Locus selection, GO analysis

The abundance of SNPs associated with the sex chromosomes made it impossible to single out any specific gene with higher association with the sex chromosome-linked plumage traits, so we compiled a list of all genes identified on the Z chromosome scaffolds, which amounted to 425 genes. Of these, 20 have been previously identified as playing a role in pigmentation, colouration, or feather development (Table 1).

**Table 1.** Table showing known or putative plumage genes located on the sapsucker Z chromosome scaffolds, what pigmentation pathway they affect, in which species that was found, and the citation for that record.

| Gene | Z scaffold | Pathway | Species | Citation |
| --- | --- | --- | --- | --- |
| RGP1 | Scaffold 1003 | melanin | *Colaptes auratus*,<br>*Setophaga coronata* | Aguillon et al. 2021,<br>Brelsford et al. 2017 |
| NNT | Scaffold 1003 | melanin | *Homo sapiens* |  |
| IDUA | Scaffold 1030 | melanin | *Homo sapiens* |  |
| ABCA1 | Scaffold 1030 | melanin | *Gallus gallus* | Attie et al. 2002 |
| XPA | Scaffold 1048 | melanin | *Homo sapiens* |  |
| FANCC | Scaffold 1073 | melanin | *Homo sapiens* |  |
| TYRP1 | Scaffold 1095 | melanin | *Coturnix japonica*,<br>*Columba livia*, *Falco*<br>*cherrug*, *Gallus gallus* |  |
| PAM | Scaffold 1030 | melanin | *Colaptes auratus*,<br>*Chrysolophus* | Aguillon et al. 2021,<br>Poelstra et al. 2015 |
| APC | Scaffold 1080 | melanin,<br>carotenoid | *Corvus corone*,<br>*Chrysolophus* | Poelstra et al. 2015,<br>Gao et al. 2018 |
| MTREX | Scaffold 1003 | melanin | *Homo sapiens* |  |
| TLN1 | Scaffold 1003 | melanin | *Homo sapiens* |  |
| SLC12A2 | Scaffold 1030 | melanin | *Homo sapiens* |  |
| MEF2C | Scaffold 1030 | melanin | *Homo sapiens* |  |
| NFIB | Scaffold 1095 | melanin | *Homo sapiens* |  |
| PTCH1 | Scaffold 1073 | melanin | *Mus musculus*, *Danio*<br>*rerio* |  |
| FST | Scaffold 1003 | carotenoid | *Erythrura gouldiae* | Toomey et al. 2018,<br>Kim et al. 2018 |
| MOCS2 | Scaffold 1004 | carotenoid | *Erythrura gouldiae* | Kim et al. 2018 |
| BMP5 | Scaffold 1044 | melanin | *Danio rerio* |  |
| LOX | Scaffold 1080 | melanin | *Danio rerio* |  |
| BNC2 | Scaffold 1082 | melanin | *Danio rerio* |  |

We identified four small autosomal regions in the LMM analyses for the non-haploblock traits (Table 2). We were able to identify three genes close to these loci. We plotted genotypes by phenotypes in mosaic plots to visualise these associations (Figure 6).

**Table 2.** Autosomal loci associated with plumage traits and the closest genes, along with Panther gene family/subfamilies.

| Scaffold | Position | Trait | Gene | Panther Family/Subfamily |
| --- | --- | --- | --- | --- |
| Scaffold 14 | 18455721 | chin | B3GLCT | Beta-1, 3-glucosyltransferase |
| Scaffold 29 | 11770566 | chin | KCNH1 | Potassium voltage-gated channel subfamily H member 1 |
| Scaffold 34 | 15391400 | throat | CD93 | Complement component C1Q receptor |
| Scaffold 34 | 15391428 | throat | CD93 |  |
| Scaffold 34 | 15391433 | throat | CD93 |  |
| Scaffold 34 | 15391989 | throat | CD93 |  |
| Scaffold 34 | 15424251 | throat | CD93 |  |
| Scaffold 172 | 166163 | black nape | no annotation |  |

**Figure 6.**
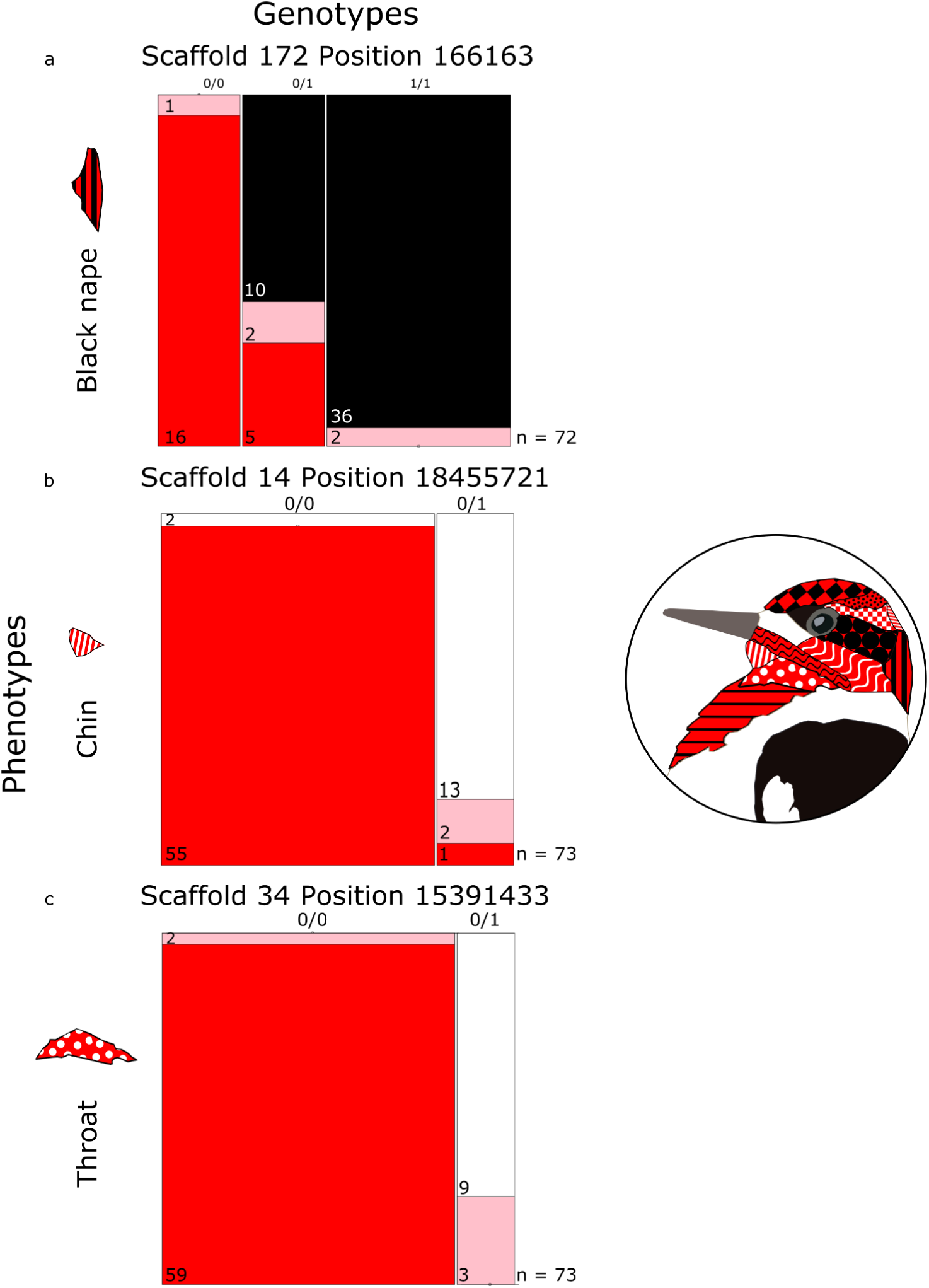
Mosaic plots of phenotype by genotype for autosomal loci linked to plumage traits in the LMM, data include all species and species crosses as well as both sexes. Red phenotypes are shown in red, white phenotypes in white, black in black, and intermediate hybrid plumages are depicted in pink for A black nape, B chin, and C throat. The number of individuals represented by each mosaic section is included on each block.

### Genotype-by-individual plots

The Z chromosome haploblock genotype-by-individual plots show long stretches of homo- and heterozygosity over vast genomic distances (on the order of tens of Mbps) (Figure 7). Typically, the plumages most common in red-breasted sapsuckers are almost exclusively homozygous and red-naped/yellow-bellied plumages are somewhat more heterozygous with many individuals homozygous for alternate alleles.

**Figure 7.**
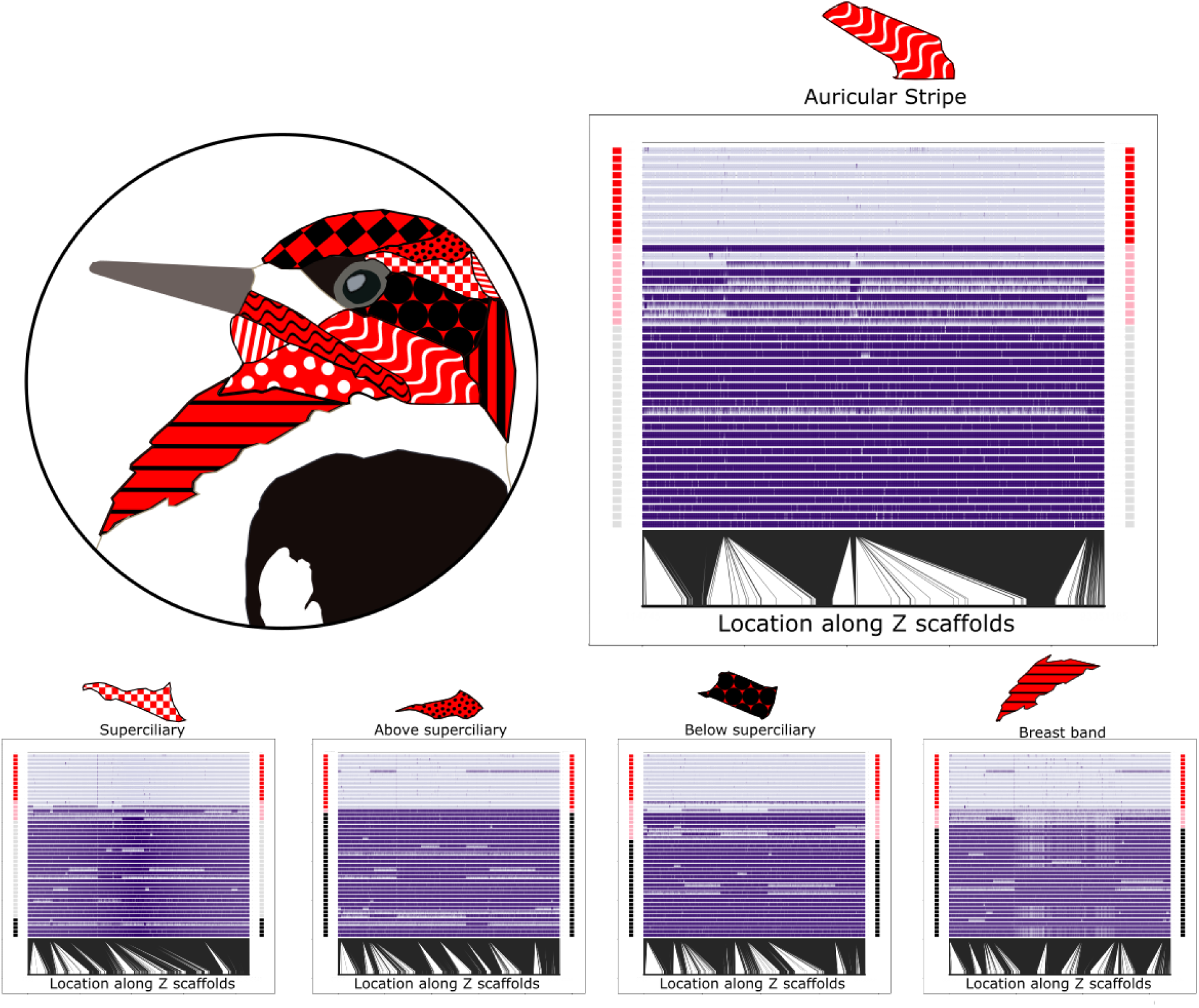
Genotype-by-individual plots of SNPs (columns) on the Z chromosome associated with each Z chromosome haploblock plumage trait for each individual (rows). Red, black, and light grey rectangles to the sides of the plot indicate the plumage trait is red, black, or white, respectively, while pink indicates an intermediate trait. Each genotype is depicted as either homozygous (purple rectangle or light purple rectangle) or heterozygous (rectangle split diagonally into a purple and light purple triangle). Lines on the bottom of the plot indicate chromosomal position of each genotype along the merged Z scaffolds.

### COG4 Locus

We searched within our reference genome for the COG4 sequence, and found it matched a region in scaffold 10 which was unannotated by liftoff. This region does not contain a high F_ST_ SNP or have associations with any plumage traits in our analyses.

## Discussion

### Red-breasted plumage is linked to large haploblocks on Z chromosome

The predominant pattern in our data indicates that the shift from the striped, sexually dichromatic plumage phenotype typical of red-naped and yellow-bellied sapsuckers to the overall red and sexually monochromatic phenotype of the red-breasted sapsucker is associated with genomic changes over large regions of the Z chromosome. As evidenced by the LMM data shown in Figure 5, -log*(p)* values on the sex chromosomes are much higher than in the rest of the genome (e.g., > 30 vs < 15). Because so many SNPs are highly associated with these plumage phenotypes, we are unable to identify any specific locus associated with the shift. Given the span of the loci affected (> 100 Mbp), it is likely that multiple genes are involved in this plumage transition. The region includes over 400 genes, 20 of which are known to have putative involvement in melanin or carotenoid pathways in vertebrates.

The red-breasted plumage probably arose fairly recently, likely in the last ∼320 thousand years, which is the estimated age of the split between red-breasted and red-naped sapsuckers (Weir & Schluter, 2004). We speculate this is the derived plumage in this trio of species for the following reasons. Because this plumage type only occurs within the red-breasted population, it likely does not result from standing genetic variation present in a shared common ancestor. Furthermore, it is common for woodpeckers to be sexually dichromatic with respect to facial striping, as seen in close relatives of these sapsuckers in particular (Williamson’s sapsuckers *Sphyrapicus thyroideus*, many *Melanerpes* spp.). Moreover, the red pigmentation in sapsucker feathers is restricted to the ends of the feather barbs, such that the bases of the feathers show the same melanin patterning across all three species Howell, 1952; Johnson & Johnson, 1985). This is consistent with head striping and dichromatic plumages as the ancestral condition in this trio and that an overall increase in carotenoid deposition, not involving melanin pigmentation, is involved in the shift. It is interesting that the loci involved in this change seem to be in areas of high F_ST_ among all three sapsucker species in this study (see Natola & Irwin, 2025). This means that it is not only the different-looking red-breasted sapsucker that is divergent in this region—rather, massive genomic shifts have occurred between the red-naped and yellow-bellied sapsuckers as well, but interestingly this has not resulted in major changes in plumage colouration between these two forms.

The genotype-by-individual plots of the LMM-identified sex chromosome-linked plumage traits (Z haploblock traits) show strong patterns of extended Z-chromosome homozygosity within each of the most differing plumage types: sapsuckers with a red trait are homozygous for one allele, and those with black or white traits tend to be mostly homozygous for the opposite allele. In general, phenotypically intermediate birds tend to show higher heterozygosity. Interestingly, this tends to occur in large haplotype blocks across the chromosome, but there does appear to be evidence of past recombination events, indicated by the switching between long regions of homozygosity and long regions of heterozygosity across the Z chromosomes. The presence of large stretches of heterozygosity in intermediate birds suggests the possible involvement of a large structural variant, likely an inversion (Todesco et al., 2020). However, the recombination, though rare, complicates this inference because an inversion would likely preclude such crossing over events like those evidenced in our broken haploblocks (Kirkpatrick, 2010).

Another observation from these data is that the red phenotype associated alleles are almost universally homozygous, meaning birds with heterozygous genotypes are almost never fully red, whereas more birds with short heterozygous stretches tend to be black or white instead of intermediate in colour. Possibly, this indicates some dominance effects. The phenotypic transition from striped heads to all red heads may be controlled by a single locus within the large linkage block, or a polygenic trait involving multiple genes within the block. The data suggest that black and white traits are dominant to red traits, and homozygosity is required for most red plumages. This contrasts to warbler examples in which the carotenoid deposition is dominant. In golden-winged and blue-winged warblers, black throat traits are recessive and white/yellow pigmented traits are dominant (Baiz et al., 2020). Similarly, a black mask in Townsend’s warblers appears to be recessive to yellow plumage in hermit warblers (*Setophaga townsendi* and *S. occidentalis,* Rohwer & Wood, 1998; Wang et al., 2020). However, in canaries red plumage is encoded by a dominant allele at one locus and a recessive allele at a second locus (Lopes et al., 2016), although this is not a natural variant, but one created through artificial selection.

### Monomorphism transition

One of the most intriguing aspects of the change to an overall red phenotype in the red-breasted sapsucker is that it is accompanied by a transition from sexual dichromatism to sexual monochromatism. Sexual dichromatism is often interpreted to result from sexually antagonistic selection, whereby natural selection drives females to be cryptic while sexual selection drives males to be more conspicuous (Gazda et al., 2020; Lande, 1980; Price & Birch, 1996). Dichromatism is thought to be a generally evolutionarily constrained trait (Lande, 1980), though over 150 transitions between dichromatism and monochromatism (in either direction) have been documented in Passerines including 130 transitions within genera, so rapid transitions such as that seen here are certainly possible (Price & Birch, 1996). When sexual chromatism does change, it appears that males and females are equally likely to drive the shift (Price & Birch, 1996).

As much as we know about examples of dichromatism and its potential evolutionary drivers, we have surprisingly few examples in which we understand the genetic and biochemical mechanisms that allow for plumage to differentiate between sexes. In some Passerine species plumage dichromatism is apparently controlled by testosterone (Sin et al., 2020; Khalil et al. 2020). In Northern Cardinals (*Cardinalis cardinalis*) male and female plumage differences are associated with *cis*-regulatory transcription factors upstream of a well-characterised pigmentation gene, CYP2J19 (Sin et al., 2020). In canaries, dimorphism is achieved through degradation of pigment produced by the gene BCO2 in female integumentary tissues. Reproductively senescent and ovariectomised female canaries have plumage similar to males, indicating the degradation of pigment likely occurs in the presence of estrogen (Gazda et al., 2020). Prior research has shown woodpecker plumage is unchanged by ovariectomy, castration, or injection of follicular hormones (Howell, 1952), so the mechanism may be different in sapsuckers and finches. Our data suggest the sex chromosomes and/or novel structural rearrangements may be places to look for transcription factors in the future to further explain the mechanism controlling the shift from di-to mono-chromatic plumage in sapsuckers.

### Autosomal associated loci

We also found autosomal loci associated with several traits (black nape, chin, throat, Figures 2, 3, 4, 6, Table 1). The chin is associated with two SNPs on separate scaffolds, scaffold 14 and scaffold 29, which are upstream of the B3GLCT and KCNH1 genes respectively. B3GLCT1 is involved in protein modification and glycosylation (Sato et al., 2006), while KCNH1 is implicated in the formation of voltage-gated potassium channels (Occhiodoro et al., 1998). The loci associated with throat pigmentation are all downstream of the CD93 gene on scaffold 34, which plays a role in cell-cell adhesion and host defense (Langenkamp et al., 2015). It is not clear how any of these cellular processes might relate to plumage. However, it is interesting that the chin and throat loci exhibit a similar pattern, wherein red plumages are homozygous for one allele, white plumages tend to be heterozygous, intermediates can be either of these genotypes, but no homozygotes for the alternate allele exist. This may be evidence that the alternate homozygous genotype is lethal, or that due to low population frequencies by chance we did not sample 1/1 homozygotes. However, these loci may also be paralogs to loci on the W chromosome. The chin of male red-naped and yellow-bellied sapsuckers is red while that of females is white, and the throat of males of both species and female red-naped is red throat while that of female yellow-bellied is white. These traits may therefore have sex linkage. It is possible that white traits appear as heterozygotes (occurring in females only) and red traits as homozygotes if a homozygous locus at the location identified is paired with a paralog on the W chromosome causing white plumage.

The black nape trait has high association with a SNP on scaffold 172, which does not map to any gene in our annotation (Table 2). This scaffold maps to the Chromosome 11 random region of the Zebra Finch assembly (RefSeq assembly accession GCF_000151805.1, Warren et al., 2010), so this part of the genome is likely poorly characterised in birds. Red and black genotypes are typically homozygous at this location for either allele, and there are no black 0/0 genotypes or red 1/1 genotypes, but heterozygotes may have any phenotype, and intermediate phenotypes may have any genotype (Figure 6).

### Plumage transition in context of sapsucker differentiation

Natola and Irwin (2025) found that sapsucker differentiation as measured by F_ST_ was high in many regions along much of the genome, especially in comparisons involving the yellow-bellied sapsucker and in regions showing high linkage disequilibrium, but that it was highest in the Z chromosomes where differentiation was extreme given the short time since divergence from their most recent common ancestor. The major transition from striped, sexually dichromatic plumage types to red, sexually monochromatic plumage appears to primarily involve differentiation on the Z chromosome. In some species, such as golden-winged and blue-winged warblers (*Vermivora chrysoptera* and *V. cyanoptera*), genomic regions associated with plumage seem to be the primary regions of diversification (Toews et al., 2016). In other systems, loci controlling plumage colouration occur within many differentiating genomic regions (Harris et al., 2018). For sapsuckers, these trends appear to occur somewhere in between wherein many other loci are differentiating, but the distinct phenotypes are associated with the most consequential of the genomic shifts.

### Selective consequences of this transition

There are several potential consequences associated with the plumage transition from striped, sexually dichromatic plumage types to red, sexually monochromatic forms among sapsuckers. For one, the bright red plumage may attract predators with good colour vision (accipiters, Walters et al., 2002a, 2002b). Conversely, the white plumage of the red-naped and yellow-bellied sapsuckers may be more conspicuous to predators that see mainly in black and white, such as their major mammalian predators (squirrels, weasels, bears, Walters et al., 2002a, 2002b). The environments in which the species live may modulate the relative importance of these effects. Gloger’s rule posits that among endothermic species, individuals living in darker and more humid habitats tend to be more heavily pigmented, and it is certainly the case that the most heavily pigmented sapsucker (more red pigment), the red-breasted, persists in the most moist climates (Walters et al., 2002a, 2002b). The monomorphism of this plumage type suggests it might not be disadvantageous for a female to also be brightly coloured in the darker forests of the west coast. Or it could suggest that a mechanism to downregulate red expression in females has not yet evolved (Price & Birch, 1996) and that a balance between natural and sexual selection in the sexual antagonism arms race is not yet resolved. Because we know that the plumage colour change is likely associated with the Z chromosome, we especially suspect that sexual antagonism is at play (Albert & Otto, 2005). No data exist on the extent of selection on either phenotype, so we can only conjecture as to the full fitness effects of either plumage type.

It may be true that this transition has effects on sexual selection as well. Sexual selection is usually invoked when species are sexually dimorphic or dichromatic (Gazda et al., 2020; Lande, 1980; Price & Birch, 1996). If there were no mating advantage to being more conspicuous, natural selection should keep males as well camouflaged as females and plumages should be sexually monochromatic. Woodpeckers tend to be sexually dichromatic, so it is likely that the differences in plumage between males and females serve as sexual signals (Howell, 1952; Short, 1972). The closest relative to these birds, the Williamson’s sapsucker, are so sexually dichromatic that males and females were initially classified as separate species (Cassin, 1852; Coues, 1874; Gyug et al., 2020; Henshaw, 1875; Newberry, 1857). We therefore believe there was strong sexual selection on ancestral sapsucker plumage types. However, sexual selection can be unspecific, wherein rather than females preferring a specific plumage trait, birds often show a preference for novelty or extreme phenotypes. Examples of this have been described in house finches (*Haemorhous mexicanus,* Hill, 1990), Java munias (*Lonchura leucogastroides*, Plenge et al., 2000), and collared flycatchers (*Ficedula albicollis*, Qvarnström et al., 2004) which prefer extreme red plumage; preference for artificially added ornaments such as elongated tails in long-tailed widowbirds (*Euplectes progne,* Andersson, 1982); or preference for completely novel artificial ornaments such as red leg bands or artificial white crest feathers in zebra finches (*Taeniopygia guttata,* Burley et al., 1982; Burley & Symanski, 1998). It is therefore possible that the extreme red of the red-breasted sapsucker plumage could be an attractant to all sapsucker females. Howell (1952) noted that in interspecific sapsucker pairings involving red-breasted and red-naped sapsuckers, the male was almost invariably redder than the female. Johnson and Johnson (1985) subsequently examined their data and found very strong patterns corroborating this finding. Natola et al. (2022) did not find evidence of this, however.

These effects are particularly relevant because assortative mating based on plumage colouration is suspected to act as a pre-zygotic reproductive barrier in many birds (Sætre et al., 1997). If a change in plumage can arise quickly and act as an isolating barrier, it could potentially accelerate diversification. However, if the novel red plumage is actually attractive to all species, as postulated by Howell (1952) and Johnson and Johnson (1985), it may promote hybridization and drive an increase in red-breasted sapsucker genotypes and swamping of red-naped and yellow-bellied genotypes. We note that hybridization between red-beasted and red-naped sapsuckers is more limited than that between red-naped and yellow-bellied sapsuckers (Howell, 1952; Johnson & Johnson, 1985; Natola et al., 2021; Seneviratne et al., 2016), the two forms that look most alike, though they are not genetically closest, and the introgression of redness from red-breasted to the other forms may be somewhat impeded.

### Importance of the Z chromosome

Natola and Irwin (2025) added the sapsuckers to the growing list of species groups for which the Z chromosome is particularly strongly genetically differentiated among species (Irwin, 2018), and the present study adds them to the much more limited list for which admixture analysis has been performed and the phenotypic variation shown to be largely encoded on the Z chromosome. Another example is yellowhammers (*Emberiza citrinella*) and pine buntings (*E. leucocephalos*) (Nikelski et al., 2024), in which highly divergent plumage phenotypes are highly associated with two major haploblocks across a large region of the Z chromosome. Due to the unusual inheritance of the sex chromosomes compared to autosomes, there are strong theoretical reasons to expect a particularly strong role for them in genetic and phenotypic differentiation reviewed by (Ellegren, 2011; Irwin, 2018). The growing use of admixture mapping in hybrid zones should allow the identification of more such cases in the wild.

### Conclusions

Our results show a disconnect between genomic differentiation and plumage colour differentiation in sapsuckers, wherein differentiation along the majority of the genome is higher between species with the most similar phenotypes than it is between sister species with vastly different phenotypes. The most striking change in phenotype in the red-breasted sapsucker is associated with high differentiation restricted to large swathes of the Z chromosome. Red carotenoid pigmentation in multiple plumage patches is associated with similar genes and regions across the Z chromosome. This large haploblock is also likely implicated in the recent transition from sexual dichromatism to sexual monochromatism within the red-breasted sapsucker. This likely has major implications for both natural and sexual selection, and on reproductive isolation among sapsuckers. Massive changes can be wrought across chromosomes over short evolutionary time spans, while coincident genomic differentiation in the same regions in other lineages may lead to very little differentiation at all.

## Acknowledgments

We are grateful to Sampath Seneviratne for obtaining DNA and photographs from many of the red-breasted hybrids used in this analysis (and analyzed previously in Seneviratne et al. 2012 and Seneviratne et al. 2016). Carla Cicero (Museum of Vertebrate Zoology), Donald McAlpine (New Brunswick Museum), Ray Poulin (Royal Saskatchewan Museum), Garth Spellman (Denver Museum of Natural Sciences), Paul Sweet (American Museum of Natural History), Ildiko Szabo (Beaty Biodiversity Museum), and Kevin Winker (University of Alaska Museum) kindly provided photographs and tissues for analysis. Kathy Martin, Loren Rieseberg, Dolph Schluter, and Anne Yoder provided helpful feedback on the manuscript. For research funding, we thank the Natural Sciences and Engineering Research Council of Canada (RGPIN- 2017- 03919, RGPAS- 2017- 507830, RGPIN- 2023- 04300) and the Linnean Society.

## CRediT Statement

LN: Conceptualization, Methodology, Formal Analysis, Investigation, Data Curation, Writing - Original Draft, Writing - Review & Editing, Visualization, Project Administration, Funding Acquisition. JH: Conceptualization, Methodology, Investigation, Writing - Review & Editing. DI: Conceptualization, Methodology, Writing - Original Draft, Writing - Review & Editing, Visualization, Supervision, Project Administration, Funding Acquisition.

## Data Availability

All raw reads are accessioned in SRA (PRJNA1295570), and all analysis scripts and metadata files will be archived in dryad upon acceptance of publication.

