## Supplemental Materials for "Admixture mapping among three hybridizing sapsucker (*Sphyrapicus*) species shows importance of the Z chromosome in phenotypic differentiation and speciation"

**Bayesian Sparse Linear Mixed Models**

In addition to Linear Mixed Models, we ran Bayesian Sparse Linear Mixed Models (BSLMM) (Zhou et al. 2013) because this method does not assume every associated locus is involved in a trait. Rather, it identifies those SNPs with the highest likelihood of involvement in the trait (Pallares et al. 2014). We ran each trait with 5 million burn-in steps and 20 million MCMC steps recording every 1,000 iterations as in Delmore et al. (2016). These yielded a Posterior Inclusion Probability (PIP), which is the frequency a variant is estimated to have a sparse effect in the MCMC, and an effect size demonstrating the magnitude of the plumage change associated with each locus. We visualized all GEMMA results with Manhattan plots made in the qqman R package.

The BSLMM models run on the non-haploblock linked traits identified a number of loci associated with plumage and with a substantial effect size (Figure S1). Effect sizes show the inferred effect each SNP has on the trait, or the magnitude of trait association with the trait (Zhou 2016). Because our traits are encoded as scores instead of direct measurements, the same plumage score can indicate different amounts of red/white/black coloration, and the exact effect sizes should be interpreted with caution. Instead, they are more useful in differentiating traits dictated by few loci of large effect as opposed to many loci of small effect.

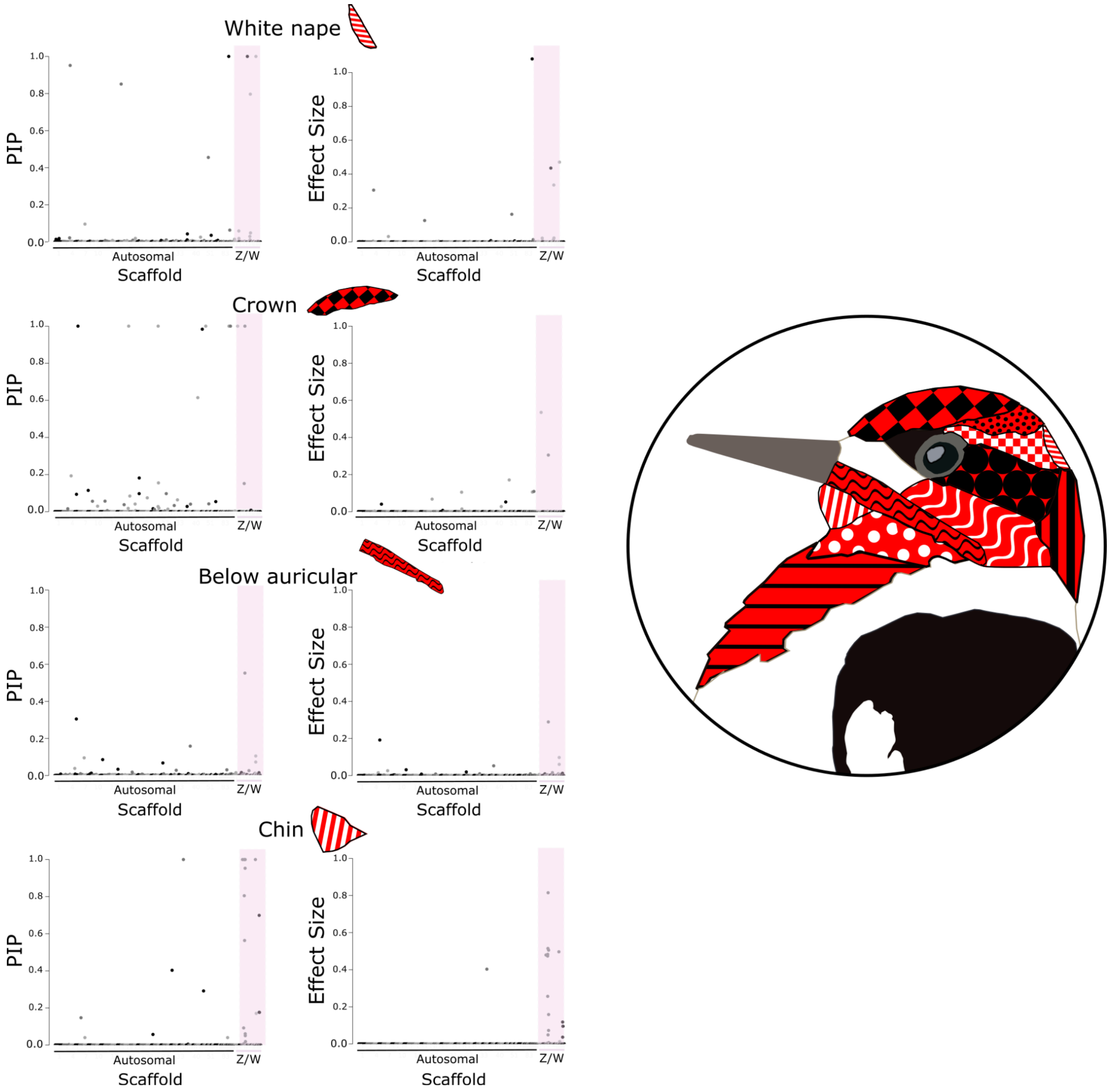


**Figure S1 Plots showing strength of association between each SNP and a phenotypic trait, as estimated using BSLMM output from GEMMA on non-haploblock traits, data include all species and species crosses as well as both sexes. Plots included show (left) Posterior Inclusion Probability (PIP), and (right) effect size, for (a) white nape, (b) crown, (c) below auricular, (d) chin. Sex chromosomes are indicated by violet shading.**
